# BOLD response to multiple grip forces in MS: going beyond the main effect of movement in BA 4a and BA 4p

**DOI:** 10.1101/2020.10.11.334599

**Authors:** Adnan A.S. Alahmadi, Matteo Pardini, Rebecca S Samson, Egidio D’Angelo, Karl J Friston, Ahmed T Toosy, Claudia A.M. Gandini Wheeler-Kingshott

## Abstract

This study highlights the importance of looking beyond the main effect of movement to study alterations in functional response in the presence of central nervous system pathologies such as multiple sclerosis (MS). Data show that MS selectively affects regional BOLD (Blood Oxygenation Level Dependent) responses to variable grip forces (GF). It is known that the anterior and posterior BA 4 areas (BA 4a and BA 4p) are anatomically and functionally distinct. It has also been shown in Healthy volunteers that there are linear (1^st^ order, typical of BA 4a) and non-linear (2^nd^-4^th^ order, typical of BA 4p) BOLD responses to different levels of GF applied during a dynamic motor paradigm. After modelling the BOLD response with a polynomial expansion of the applied GFs, the particular case of BA 4a and BA 4p were investigated in Healthy Volunteers (HV) and MS subjects. The main effect of movement (0th order) analysis showed that the BOLD signal is greater in MS compared to healthy volunteers within both BA 4 sub-regions. At higher order, BOLD-GF responses were similar in BA 4a but showed a marked alteration in BA 4p of MS subjects, with those with greatest disability showing the greatest deviations from the healthy response profile. Therefore, the different behaviour in HV and MS could only be uncovered through a polynomial analysis looking beyond the main effect of movement into the two BA 4 sub-regions. Future studies will investigate the source of this pathophysiology, combining the present fMRI paradigm with blood perfusion and non-linear neuronal response analysis.

## 1. Introduction

The primary motor cortex, M1 or Brodmann area 4 (BA 4) is very important because of its essential role in generating movement, a skill often affected by diseases such as Multiple Sclerosis (MS). Interestingly, BA 4 has two sub-regions with distinct cytoarchitectonic properties, anatomy, and neurochemistry both in humans and primates (Strick and Preston, 1982, Geyer et al., 1996, Kaas and Collins, 2002). Geyer et al showed that the two sub-divisions have differences in transmitter binding sites and laminar density of neurons (Geyer et al. 1996). In particular, they showed that BA 4a has more densely packed pyramidal cells. On the other hand, BA 4p has higher laminar specific densities of different receptor and transmitter binding sites.

Magnetic Resonance Imaging (MRI) functional studies have shown that in healthy volunteers the blood oxygen level dependent (BOLD) signal response is modulated by attention (Binkofski et al., 2002) and imagined forces (Sharma et al., 2008) in BA 4p, whilst BA 4a responds to motor control (Alahmadi et al., 2016). In other words, BA 4a is predominantly related to execution whereas BA 4p is predominantly related to higher-order cognitive tasks (Sharma et al., 2008, Binkofski et al., 2002, Alahmadi, 2020, Alahmadi et al., 2015b, Alahmadi et al., 2017, Alahmadi et al., 2016).

A recent study of healthy volunteers has revealed that the BOLD response to complex motor tasks, involving different grip forces (GFs), is characterised by different MRI signal response profiles, even during observation (Alahmadi et al., 2016, Casiraghi et al., 2019, Keisker et al., 2009). Interestingly, the study reported a distinct behaviour of the BOLD-GF relationship within the two sub-regions of BA 4 (Alahmadi et al., 2016). The BOLD-GF relationship follows a distinct non-linear negative third order profile within BA 4p, while it is linear in BA 4a. Moreover, in healthy volunteers the BOLD signal within the two sub-regions has a distinct response to motor complexity, when using the dominant or non-dominant hand while applying different GFs (Alahmadi et al., 2015b). Therefore, the functional differences between BA 4a and BA 4p provide an ideal case to compare complex BOLD responses going beyond the 0-order (or main effect) of movement in a pathology like MS.

A question arises as to whether and if so how, these behaviours are affected by aging and by diseases of the central nervous system, warranting a full characterization of the BOLD behaviour in these two sub-regions, beyond a standard main effect of movement. Some investigations have reported, indirectly, that there are distinct responses in BA 4p and BA 4a in aging (Ward and Frackowiak, 2003) and in stroke patients (Ward et al., 2007). In particular, they showed that BA 4p is affected by aging and that it is key to the functional integrity of the cortical-spinal system and motor recovery in patients with stroke. Therefore, advanced analysis of BA 4a and BA 4p could highlight mechanisms of functional alterations otherwise shadowed in an undifferentiated analysis of BA 4.

On the bases of these considerations, the present study investigates the non-linear behaviour of the BOLD response to different GFs within BA 4a and BA 4p, in healthy volunteers and in people with MS, a neurological disease known to affect the motor system. MS has complex disease mechanisms involving a number of pathophysiological components, including demyelination, axonal loss and inflammation. Accumulation of sodium ions in tissue and a redistribution of sodium channels along damaged axons alters conduction properties (Paling et al., 2013, Cercignani et al., 2017), which could also be affected by alterations of tissue blood perfusion (Bester et al., 2015, Rashid et al., 2004, Paling et al., 2014, Rovaris et al., 2002, Lapointe et al., 2018, Rocca et al., 2007). FMRI studies in MS have shown altered patterns of activations (Rocca et al., 2007, Filippi et al., 2013, White et al., 2009) and altered resting state networks (Castellazzi et al., 2018), but were not designed to answer questions about complex BOLD behaviour.

The hypothesis of this study is therefore that, in MS, there is (1) an altered functional response in BA 4 compared to healthy volunteers during a motor fMRI task and that (2) this alteration is region-specific. Given the involvement of BA 4p in higher-order motor control and its modulation by attention and task complexity we also hypothesised that (3) area BA 4p may show more severe abnormalities than BA 4a in the presence of MS pathology when compared to healthy volunteers. If this is true, then the BOLD-GF relationship in BA 4a and BA 4p may show different regional patterns of alteration compared to Healthy volunteers, offering new insights in the pathology of MS.

## 2. Methods

### 2.1. Subjects

14 right-handed healthy volunteers (9 female, 5 male; mean age 31 (± 4.64) years) and 14 right-handed relapsing remitting MS (RRMS) patients (10 female, 4 male; mean age 35 (± 5.36) years; median (range) expanded disability status score (EDSS) 3.5 (1.5-6.5)); median (range) 9-Hole Peg Test (9-HPT) = 20.05 (14.7-33.1) were recruited. The handedness of subjects were assessed according to the Edinburgh handedness scaling questionnaire (Oldfield, 1971). All subjects gave informed consent and the study was approved by the local research and ethics committee.

### 2.2. MRI acquisition

A 3.0 T MRI scanner Philips Achieva system (Philips Healthcare, Best, The Netherlands) and a 32-channel head coil were used. The imaging acquisition protocol included the following: T1-weighted volume (3DT1): 3D inversion-recovery prepared gradient-echo (fast field echo) sequence with inversion time (TI) = 824ms, echo time (TE)/repetition time (TR) = 3.1/6.9ms, flip angle = 8° and voxel size = 1mm isotropic; BOLD sensitive T2*-weighted echo planar imaging (EPI): TE/TR = 35/2500ms, voxel size = 3×3×2.7 mm^3^, inter-slice gap of 0.3 mm, SENSE factor = 2, number of slices = 46, acquired with descending order, field of view = 192×192mm^2^, number of volumes = 200, number of dummy scans = 5, flip angle = 90**°**.

### 2.3. FMRI paradigm

The experimental design was a *visually* guided event-related fMRI paradigm, where subjects used their right (dominant) hand to squeeze a rubber ball with varying GF levels. The design comprised 5 GF targets (20, 30, 40, 50 and 60% of subjects’ maximum voluntary contraction (MVC)) interleaved with rest intervals, each repeated randomly 15 times. This paradigm has been validated previously in studies of Healthy volunteers (Alahmadi et al., 2015b, Casiraghi et al., 2019, Alahmadi et al., 2017, Alahmadi et al., 2016).

Before the fMRI session, subjects were given a 6 minute training session including learning, watching and performing a similar but not identical paradigm. During the fMRI session, participants lay supine on the scanner bed throughout the experiment and were instructed to extend both of their arms in a relaxed comfortable position. A support hand pad was provided for each subject to ensure comfort and compliance.

### 2.4. Image pre-processing and analyses

Data processing was performed using statistical parametric mapping (SPM12 (www.fil.ion.ucl.ac.uk/spm) implemented in Matlab14b (Mathworks, Sheborn, MA). The pre-processing steps for each subject adopted the following fMRI pipeline: Slice time corrections; Spatial volume realignments; (iii) Co-registration with the 3DT1 volume; (iv) Normalization with the tissue probability maps of SPM12; (v) Smoothing of the functional volumes with an 8mm isotropic full-width half maximum (FWHM) Gaussian kernel.

### 2.5. Statistical analysis

#### Within-subjects (first level analysis)

Signal changes were modelled using a polynomial expansion as described in (Alahmadi et al., 2016) and according to (Buchel et al., 1998, Buchel et al., 1996). Briefly, for each subject, a fixed effects analysis was performed. To test efficiently for linear and nonlinear effects, a parametric design was chosen. The parametric design included a set of orthogonalised polynomial orders (up to the fourth order) and was specified by the integral of the forces for each subject. This step creates five regressors of interest: 0^th^ order represents the main effect of gripping regardless of the applied force, 1^st^ order represents any linear changes between BOLD signals and the applied forces, and 2^nd^ – 4^th^ orders represent nonlinear effects modelling the relationships between BOLD signals and the applied forces. These covariates were then convolved with a canonical hemodynamic response function (HRF) for standard SPM and general linear model (GLM) analysis (Friston et al., 1995). The movement related realignment parameters were included as regressors of no interest in each GLM (Friston et al., 1996). At this within-subject level, *t*-statistics were used to test for the effects of the polynomial coefficients.

#### Between-subjects (second level analysis)

Contrast images from the within-subject analysis for the five polynomial orders were entered into random effects analyses, testing for non-linear effects within and between groups, with the appropriate *t*-tests (i.e. one sample *t*-test for the within group tests and two sample *t*-test for between group comparisons). Significant voxels were defined using *P*<0.05, corrected for multiple comparisons (FWE). Importantly, the number of comparisons (voxels) performed in this study was within the sub-regions of BA 4, thus the numbers of comparisons were low as compared to the whole brain.

In addition to the above whole brain SPM analysis, BA 4 was defined and subdivided according to (Eickhoff et al., 2005) as guided by (Geyer et al., 1996) (figure 1). The study by Eickhoff et al. 2005 introduced a new probabilistic cytoarchitectonic map of different brain regions (including BA 4), which significantly improved the accuracy of anatomical labelling. The new cytoarchitectonic map measured the probability of a single voxel falling within an area based on 10 post-mortem brains normalized to the MNI single subject template. Only voxels (or clusters) having the highest probability of falling within the MNI sub-division of BA 4 were considered for each *post mortem* subject and were used to define a map of BA 4 in MNI space. Since this is an *a priori* predefined region of interest (ROI), we used this anatomical ROI for each subject and performed small volume correction (SVC) to increase our sensitivity when looking at individual subject behaviours and parametric effects. To assess the relationship between BOLD signal and GF, we plotted the average group signal estimate over voxels within these predefined ROIs for either BA 4a or BA 4p. Each plot therefore reports the maximum likelihood estimates of the mapping between the different applied GF and BOLD signals based on the polynomial expansion (Alahmadi et al., 2016).

**Figure 1:**
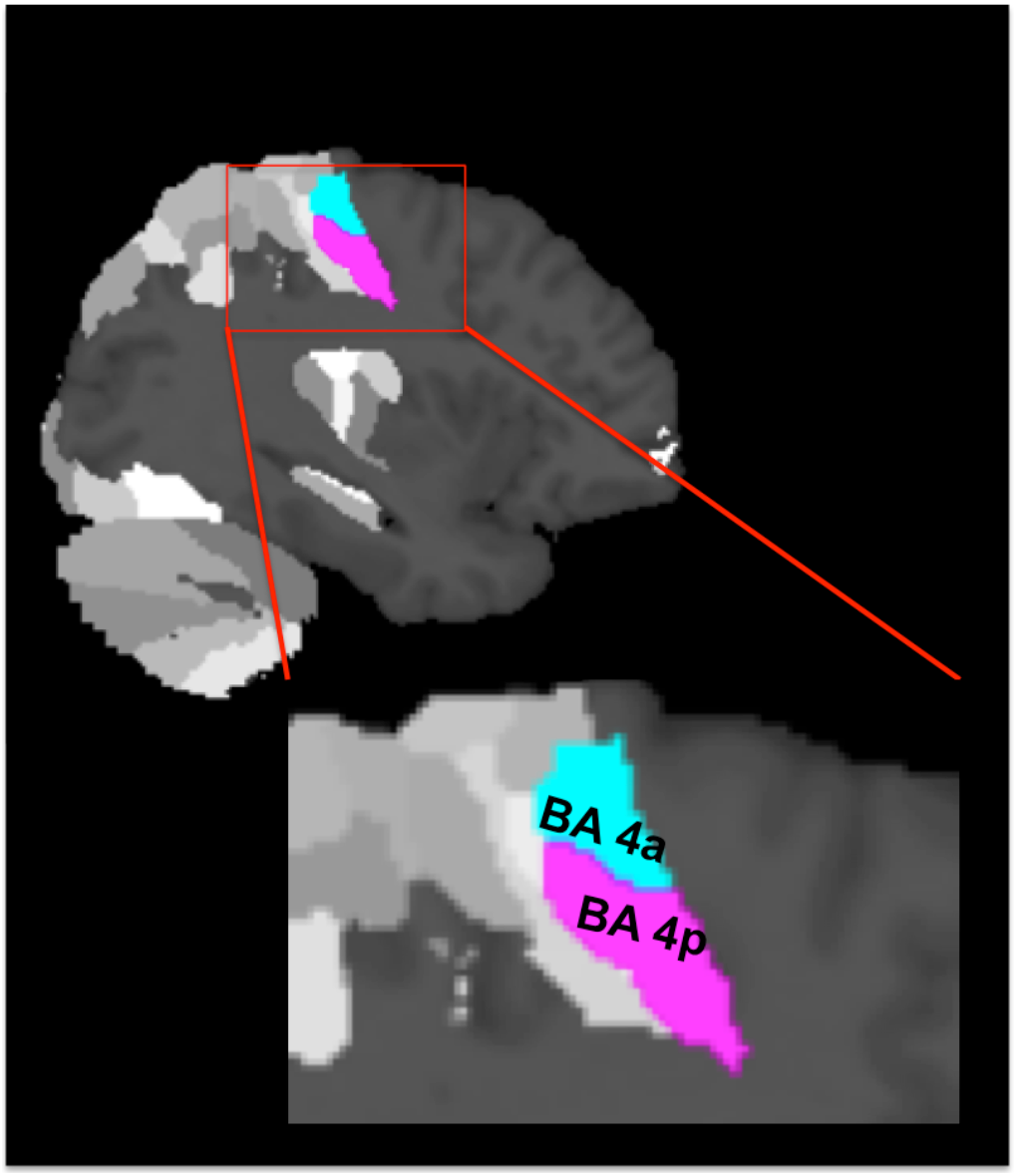
The cytoarchitectonic assignments of BA 4a and BA 4p projected onto the maximum probability map of the brain as provided by the SPM anatomy toolbox.

To better understand the effect of disability on BOLD responses to GF, we divided the MS group based on their EDSS score into two sub-groups of low (EDSS ≤ 3) and high disability (EDSS >3). The 9-HPT was highly correlated with the EDSS scores and thus the two groups could be sub-divided similarly to low (9 HPT ≤21.7) and high disability (9-HPT > 21.7). Following the sub-division of the MS group, the low EDSS group comprised 8 subjects (4 female, 4 male; mean age 33.8 (± 4.5) years; median (range) EDSS 1.5 (1.5 – 2.5); median (range) 9-Hole Peg Test (9-HPT) = 19.1 (14.7-22.1)), while the high EDSS group comprised the remaining 6 subjects (5 female, 1 male; mean age 36.1 (± 7.8) years; median (range) EDSS 6 (3.5 – 6.5); median (range) 9-Hole Peg Test (9-HPT) = 25 (22.3-33.1)).

In order to characterise the response profile of each sub-division in each group, we used the relative contribution of each polynomial term as described in Alahmadi et al. 2015b. We therefore classified the effect of GF at each voxel within BA 4a and BA 4p based on the polynomial order that showed the highest standardised group effect size (i.e. the most significant group difference).

## 3. Results

All subjects were able to perform the task correctly (table.1).

**Table 1.**
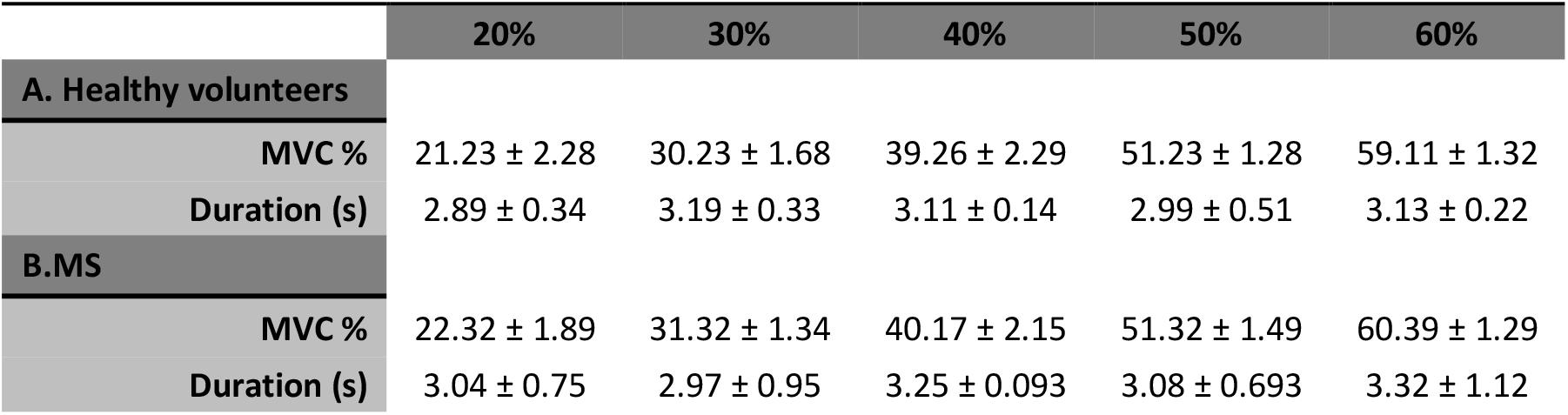
Grip force task performance, showing the average (±SD) MVC (%) and duration (s) of squeeze for healthy and MS subjects.

In this study, we report five major findings:

### 3.1. Main effect of movement

Both groups activated BA 4a and BA 4p (figure 2 and figure 3). RRMS patients showed increased and greater activation extent compared to Healthy volunteers in both BA 4a and BA 4p sub-regions (figure 2 & figure 3) (p-value=0.001). RRMS patients also showed increased activations as their EDSS increased within BA 4p only (p-value=0.001) (figure 4).

**Figure 2:**
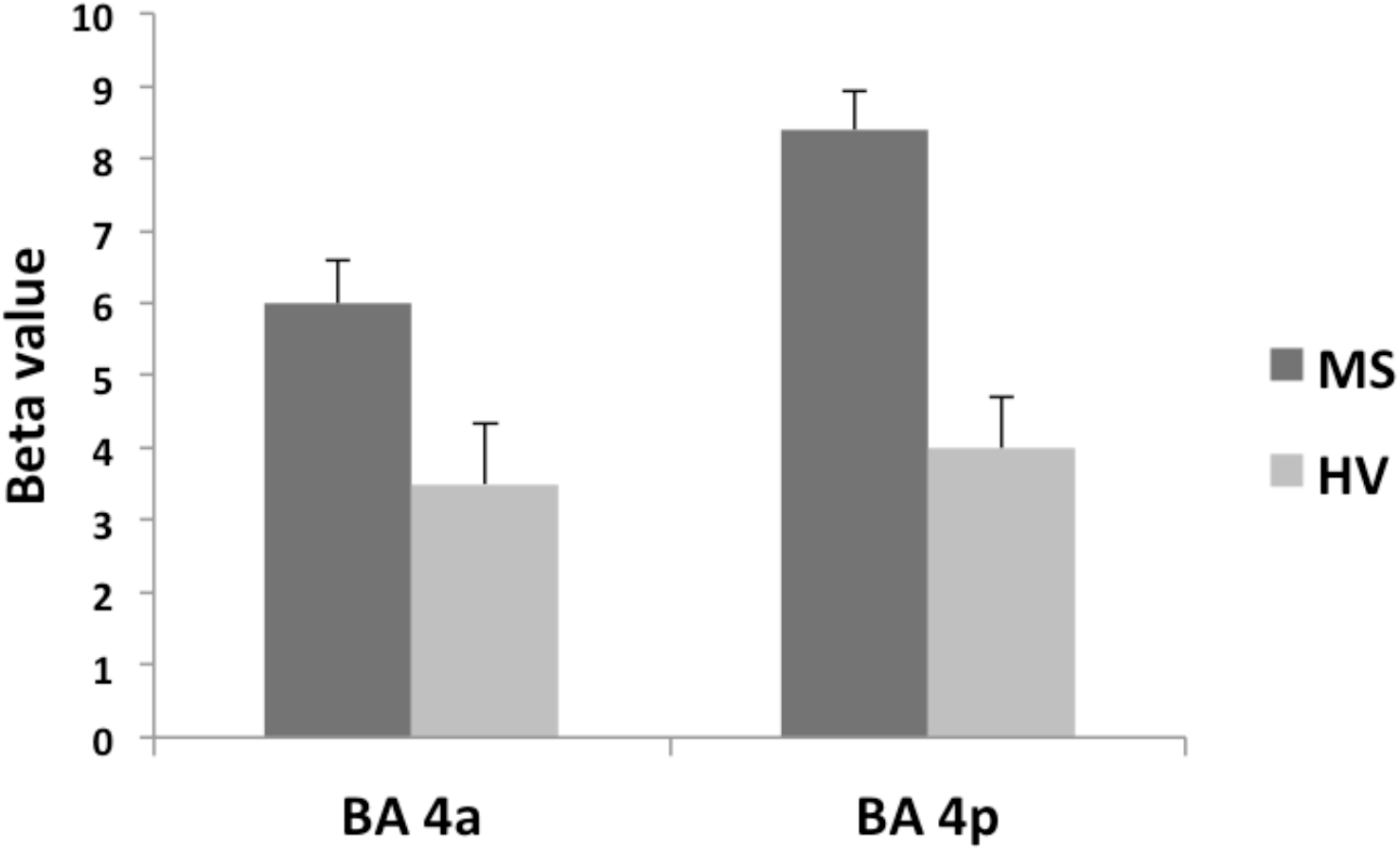
Mean of the beta values and their standard errors calculated at group level for the main effect of gripping for both groups and sub-regions. There are significantly higher betas in the MS compared to the Healthy volunteers within the sub-regions.

**Figure 3:**
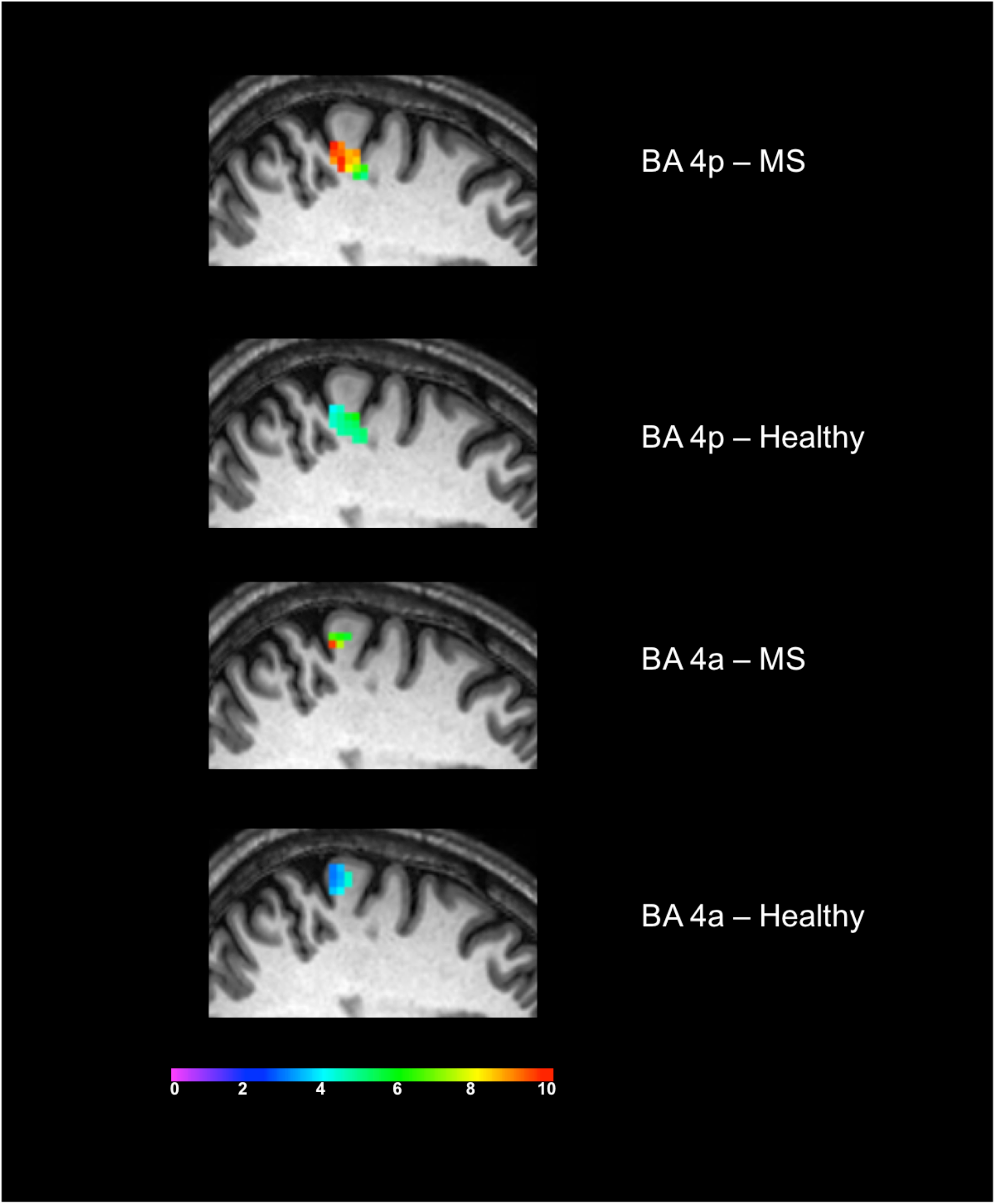
Significant activations of the main effect of gripping within BA 4p and BA 4a in both MS and Healthy volunteers. Colours represents the T-value of the effects at a 0.05 FWE.

**Figure 4:**
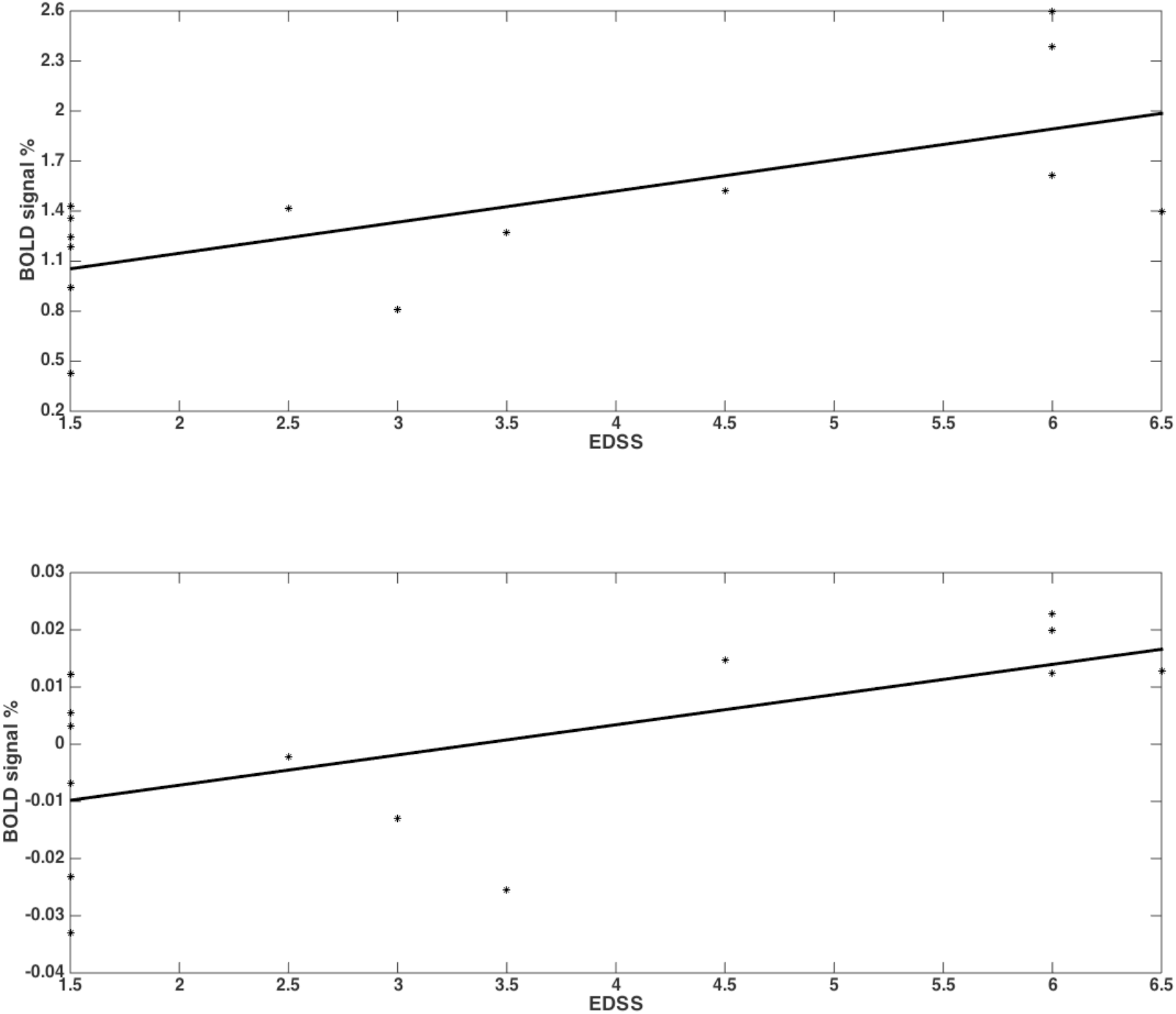
RRMS patients showed increased activations as their EDSS increased within BA 4p (p-value=0.001;r=0.68) (top plot) and within BA 4a (p-value=0.001;r=0.61) (bottom plot) in the main effect of gripping (i.e. 0^th^ order).

### 3.2. Average BOLD relationship with GF in BA 4a

In both groups, significant relationships were detected between BOLD signal and GF. There were no differences detected between MS subjects and Healthy volunteers in terms of the relationship between BOLD signal and GF within BA 4a (figure-5).

**Figure 5:**
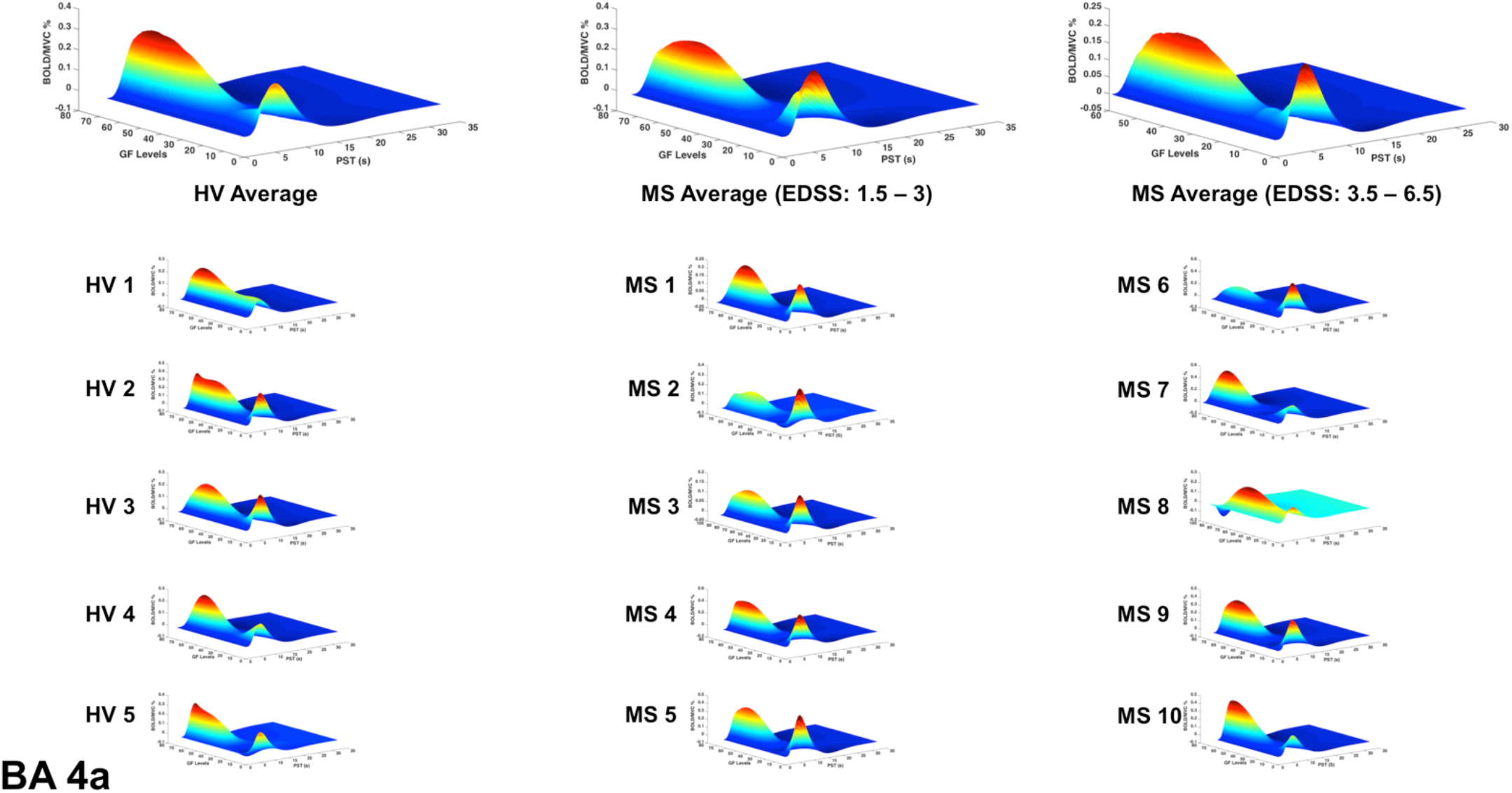
BOLD responses (Z-axis) of the fitted polynomial-orders of GF (Y-axis) at the defined post-stimulus time (PST) (X-axis) within BA 4a for Healthy volunteers (HV 1-5), MS patients with low (MS 1-5) and high EDSS (MS 6-10)—representing an estimate of the mapping between GF and BOLD based on all components of the polynomial expansion. The top row shows the average group effect while underneath examples of individual subjects are plotted.

### 3.3. Average BOLD relationship with GF in BA 4p

In patients with low EDSS, the BOLD-GF relationship was very similar to Healthy volunteers (mainly negative 3^rd^ order), whereas at higher EDSS, the predicted BOLD versus GF deviated from the Healthy volunteers pattern and the BA 4a pattern (figure 6, 3^rd^ column).

**Figure 6:**
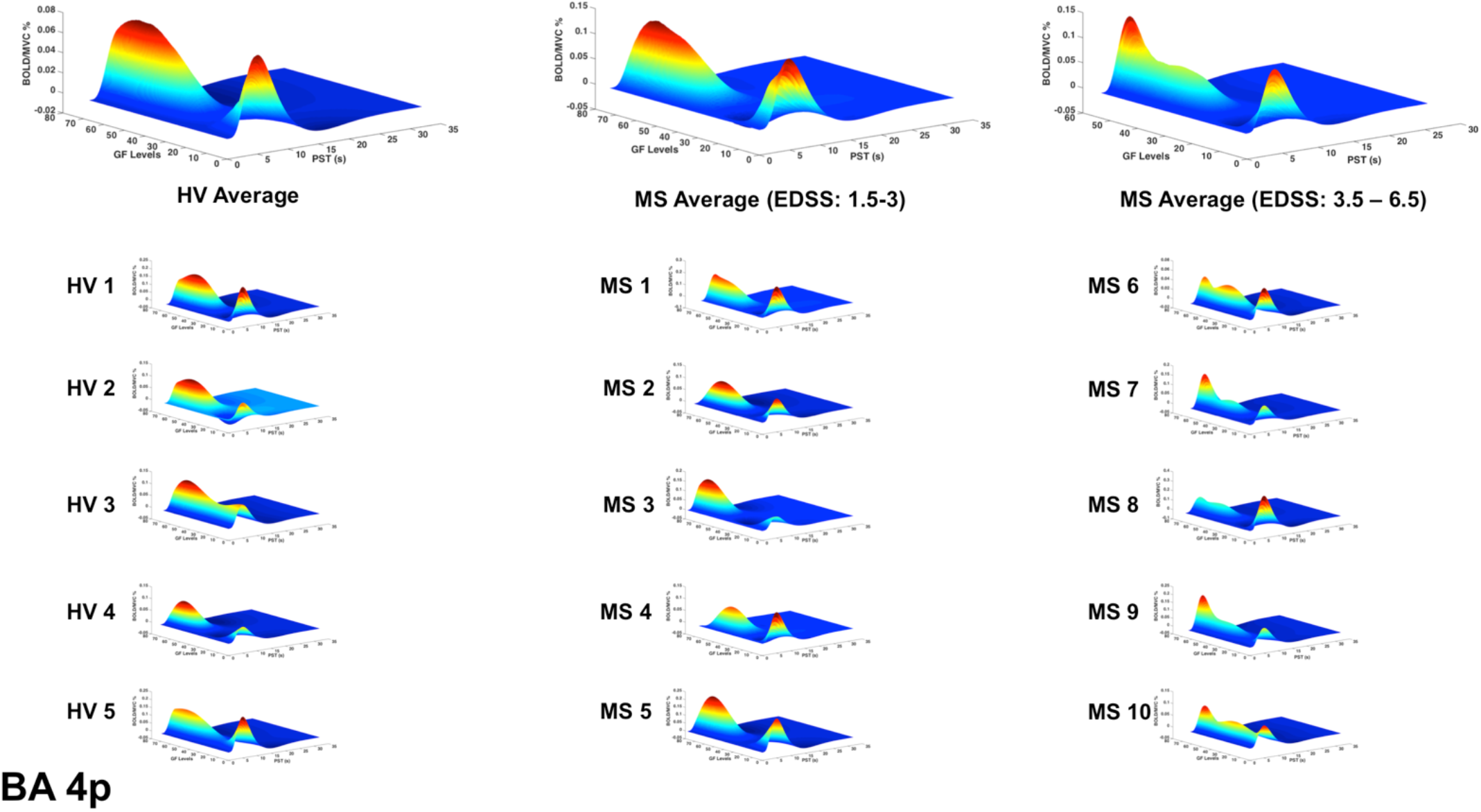
BOLD responses (Z-axis) of the fitted polynomial-orders of GF (Y-axis) at the defined post-stimulus time (PST) (X-axis) within BA 4p for Healthy volunteers (HV 1-5)), MS patients with low (MS 1-5) and high EDSS (MS 6-10)—representing an estimate of the mapping between GF and BOLD based on all components of the polynomial expansion. The top row shows the average group effect while underneath examples of individual subjects are plotted.

### 3.4. GF Response profile in single subjects

Interestingly, the profile of the mean BOLD signal versus GF was very similar across individual subjects, when grouping subjects by disease stage. Figure 5-6 show the maximum likelihood estimates of the mapping between the applied GF and BOLD signals based on the polynomial expansion at the group and subject levels.

### 3.5. Categorizing effect sizes

Performing a post-hoc analysis to identify the polynomial order showing the highest effect size, we showed that in both groups (healthy and MS) the 1^st^ order effect was predominant within BA 4a. On the other hand, the predominant effect within BA 4p was different in the two groups. In Healthy volunteers, a negative 3^rd^ order effect was predominant while in MS a positive (U-shaped) 2^nd^ order effect was predominant, with BOLD signal increasing with the highest force in MS, which elicited the lowest BOLD response in Healthy volunteers. See supplementary material.

## 4. Discussion

In this study, we demonstrate the importance of characterising high-order BOLD responses to GF as opposed to simply assessing group differences in the main effect of movement (0-order) between patients and healthy controls in pathologies such as MS. This characterisation was performed within the sub-divisions of the primary motor region (BA 4a and BA 4p), which present different order response to the intensity of movement (Alahmadi et al., 2015a, Alahmadi, 2020, Alahmadi et al., 2016). The main findings are that there are distinct grip (i.e. main effect) and force level effects within these two sub-regions in healthy and MS subjects. We further demonstrated that the main effect of gripping and the non-linear BOLD-GF relationship within BA 4p changes with disease progression.

A key finding is that the non-linear relationship between BOLD response and GF was confirmed in both Healthy volunteers and MS groups and that in people with MS it was distinctly altered only in BA 4p (figure 6). This result can be supported by the fact that the neuronal and cellular structures as well as chemistry of these two sub-regions are different (Strick and Preston, 1982, Geyer et al., 1996, Kaas and Collins, 2002). Thus, pathologies such as MS could additionally affect these two sub-regions differentially. Focusing on BA 4a and 4p enabled us to characterise the BOLD signal response based on its relationship to increasing GF, assessing its sensitivity to pathological changes at a sub-regional level.

BA 4 as a whole responded to the main effect of grip in both MS and healthy control groups. This is in line with previous motor gripping studies that showed the role of BA 4 in motor generation and function (Rocca et al., 2007, White et al., 2009, Kuhtz-Buschbeck et al., 2008, Kuhtz-Buschbeck et al., 2001, Keisker et al., 2009).

This distinct functional segregation and parametric responses was effectively captured by this study. Indeed, our findings regarding the main effect of movement show that both sub-regions are activated within both groups. The responses though are different within the two sub-regions as the BOLD signal was significantly increased in MS compared to the healthy group, especially in BA 4p (figure 2 & 3). Previous reports of motor functional studies illustrate different outcomes in the main effect of movement (within BA 4 as a whole) in MS, with the BOLD signal response shown to either increase (Lee et al., 2000, White et al., 2009, Reddy et al., 2000) or to have no difference compared to Healthy volunteers (White et al., 2009, Mancini et al., 2009). Unfortunately, most of these studies used automated anatomical labelling that relies on a template (e.g. the Talairach) or manual eye assessment labelling, both of which are highly prone to inaccuracy and difficult to generalize. The other limiting factor is that most of these studies were not anatomically specific for BA 4a or BA 4p (e.g. both M1 and S1 (primary sensory area) were labelled as “sensorimotor cortex” or SMC). This makes comparisons between our results and earlier studies difficult. The inconsistency among earlier studies can be attributed to factors such as differences in paradigms, number of subjects, threshold values, and MS subtypes. The last factor, i.e. MS subtype, may be very significant in affecting outcome as functional activations have been shown to be altered during disease progression (Rocca et al., 2005) (Rocca et al., 2005). More importantly, it has been suggested that MS patients during the early stage of the disease tend to have normal patterns of activation (Pantano et al., 2015).

This study goes beyond the main effect of movement and shows that the non-linear relationship BOLD-GFs within BA 4 is complex and region-specific. In previous studies of Healthy volunteers we have demonstrated the complexity of BOLD signals as a function of GF in different motor, sub-motor, associative and cerebellar areas (Alahmadi et al., 2015b, Alahmadi et al., 2017, Alahmadi et al., 2016, Casiraghi et al., 2019). These were seen as well in action observation and execution networks (Casiraghi et al., 2019). In the present study, we show that pathology affects the BOLD-GF relationship differently in BA 4a and BA 4p and that the altered behaviour was indicative of high EDSS. What was particularly striking was the consistency of the profile between individual subjects, reflecting the group level findings (figures 5 & 6).

The observation that the BOLD response to different GFs within BA 4p was similar to that of Healthy volunteers in patients with low EDSS, while it was consistently altered at higher EDSS poses interesting mechanistic questions, suggesting that differences not only in cytoarchitecture but also in chemoarchitecture and myeloarchitecture of these two sub-regions may translate into differences in their susceptibility to MS pathology. For example, these architecture properties as well as differences in the distribution of neural cells densities within the two sub-regions (Geyer et al., 1996) could be altered or be the cause of alterations in myelination, axonal loss, vascular or neuronal activity in MS. One could speculate that given the rich density of neurotransmitters in BA 4p compared to BA 4a (Geyer et al., 1996), our observations could reflect an impaired neuronal response. With the present data, though, it is not possible to link the present functional findings to tissue microstructure alterations, nor to infer a causal relationship between an impaired functional response to a complex task and blood perfusion, microvascular response, or even sodium channels malfunction, all of which are known to be regionally affected and potentially responsible for our observations. Furthermore, these differences could be due to differences in their structural connectivity to other brain regions that could potentially drive this different behaviour. Thus, our findings underline the need for multi-modal and longitudinal studies that could include other quantitative techniques such as diffusion weighted imaging (DWI) (to assess tissue microstructure and connectivity), MR spectroscopy (to assess metabolic changes), sodium imaging (to assess effects of sodium ions tissue distribution essential for neurotransmission), grey matter sensitive sequences (to assess grey matter atrophy and lesions) and perfusion imaging to pin down mechanistic hypothesis and deliver sensitive and specific *in vivo* imaging biomarkers of the functional substrate of MS alterations; we believe that it is really important to go beyond reporting the main effect of movement, i.e a change in BOLD signal amplitude.

The fact that these behaviours reflected EDSS association is also of interest. Previous studies have suggested that changes in functional activations in MS are possibly related to compensatory mechanisms (Staffen et al., 2002, Audoin et al., 2003, Lenzi et al., 2007, Mainero et al., 2004b), which could also be advocated to explain the higher activations observed in the main effect of gripping in MS compared to healthy volunteers. It should be noted, however, that previous studies showed that BA 4p is involved in executive motor function tasks compared to BA 4a. The predominant reported factors for BA 4p involvement were attention (Binkofski et al., 2002), complexity (Alahmadi et al., 2016) and imagination (Sharma et al., 2008). In the current study, these factors, especially attention and complexity, are all invoked by task execution. The use of an increased GF increases the complexity of performance and the use of visual feedback to reach (as quickly as possible) and maintain (as precisely as possible) forces at a specific level (especially low and high GFs) requires increased attention. Previous reports showing that in MS there are functional alterations with task complexity (Filippi et al., 2002) and attention (Mainero et al., 2004a), may support our finding of increased functional changes in BA 4p. A positive correlation of the BOLD effect with EDSS, though, indicates that this increased activity may either be a failed compensatory mechanism or may not be compensatory after all, but rather maladaptive. Advanced network modeling of the functional signal behavior may assist in understanding the source of these alterations (Friston et al., 2003). Dynamic causal modeling (DCM) would be a possible analysis to perform to help understand the precise nature of the non-linearity in the BOLD response at the neuronal or hemodynamic level. Given that the most interesting hemodynamic nonlinearities emerge over a timescale of seconds, due to hemodynamic saturation effects (Friston et al., 2003, Friston et al., 1996, Friston et al., 1998, Friston et al., 2000), a revised paradigm that includes variations in the GF duration as well as strength would be desirable for future experiments.

### Limitations and methodological considerations

In this study, there are some methodological considerations to disclose. The number of subjects could be considered to be relatively low. This is especially relevant when sub-dividing the MS subjects into two different groups. However, the subdivision of patients into two groups is exploratory and the study was not powered for it. Given the striking results, though, it is important to report these findings, which could drive future larger studies. Another limitation of this study is the inability to investigate lesions within the targeted ROIs (i.e. BA 4). Grey matter lesions in MS have been reported using MRI (Sethi et al., 2012, Geurts et al., 2011). However, grey matter lesion sequences were not planned here, although the authors advocate the need to include a grey matter lesion sequence in future functional MRI studies of MS.

## 5. Conclusion

BA 4 has two sub-divisions (BA 4p and BA 4a) that are anatomically and functionally distinct; therefore, BA 4 lends itself very well to the investigation of sub-regional differences in functional response to complex motor tasks in healthy and MS subjects. Here, we demonstrated region-specific alternations in the BOLD response to movement, showing that we should investigate beyond the main effect to unveil altered non-linear coupling between the BOLD signals and GFs, especially within BA 4p. The information obtained from studying the BOLD-GF relationship could reflect not only disease activity, but also the degree of morbidity. Furthermore, the alteration of the BOLD-GF profile in high EDSS patients compared to Healthy volunteers is a very interesting finding that opens an entire new set of avenues to study the mechanisms of neurological diseases *in vivo*. For example, the consistent alteration of the BOLD-GF profile in BA 4p in the advanced stage of MS could be explained by many factors: perhaps demyelination of BA 4p itself, which could make it unable to support an efficient functional response? Or perhaps white matter fibers subtending BA 4p have redistributed sodium channels that – instead of supporting functionality – impair efficient neurotransmission? Or perhaps in MS the microvascular response is impaired with a regional specificity that makes BA 4p responding differently than BA 4a? These questions highlight the need for multi-modal cross-sectional and longitudinal studies that aim at disentangling the contribution to functional alterations of many factors, in order to pin down sensitive and specific *in vivo* imaging biomarkers.

## 6. Acknowledgements

CGWK receives funds from the UK MS Society (#77), Wings for Life (#169111), Horizon2020 (CDS-QUAMRI, #634541), BRC (#BRC704/CAP/CGW); AA was supported by KAU. MP was supported by the AKWO association, Lavagna (Italy). KJF is supported by the Wellcome trust. RS also receives funding from the UK MS Society (#77) and the NMR Research Unit, Queen Square MS Centre receives support from the UCL-UCLH Biomedical Research Centre. ED was supported by the European Union’s Horizon 2020 Framework Programme for Research and Innovation under Specific Grant Agreement 720270 (Human Brain Project SGA1) and 785907 (Human Brain Project SGA2), with specific involvement of the CDP2 subproject. The work was also sponsored by the MNL project of the Centro Fermi (Rome, Italy). ATT is supported by grants from Rosetrees Trust (MS632), UK MS Society (984) and Medical Research Council (MR/S026088/1).

## Notes

### Competing Interest Statement

The authors have declared no competing interest.

